# Benzene metabolites increase vascular permeability by activating heat shock proteins and Rho GTPases

**DOI:** 10.1101/2024.12.04.626801

**Authors:** Igor N. Zelko, Ahtesham Hussain, Marina V. Malovichko, Nalinie Wickramasinghe, Sanjay Srivastava

## Abstract

Benzene is a ubiquitous environmental and occupational pollutant abundant in household products, petrochemicals, and cigarette smoke. It is also a well-known carcinogen and hematopoietic toxin. Population-based studies indicate an increased risk of heart failure in subjects exposed to inhaled benzene, which coincides with the infiltration of immune cells into the myocardium. However, the mechanisms of benzene-induced cardiovascular disease remain unknown. Our data suggests that benzene metabolites trans,trans-muconaldehyde (MA), and hydroquinone (HQ) propagate endothelial activation and apoptosis analyzed by endothelial-specific microparticles in C57BL/6J mice plasma. Subcutaneous injections of MA and HQ increased vascular permeability by 1.54 fold and 1.27 fold correspondingly. In addition, the exposure of primary cardiac microvascular endothelial cells to MA increased vascular permeability detected by transendothelial monolayer resistance and by fluorescently labeled dextrans diffusion. The bulk RNA sequencing of endothelial cells exposed to MA for 2, 6, and 24 hours showed MA-dependent upregulation of heat shock-related pathways at 2 and 6 hours, dysregulation of GTPases at 6 hours, and altered cytoskeleton organization at 24 hours of exposure. We found that the HSP70 protein induced by MA in endothelial cells is colocalized with F-actin foci. HSP70 inhibitor 17AAG and HSP90 inhibitor JG98 attenuated MA-induced endothelial permeability, while HSP activator TRC enhanced endothelial leakage. Moreover, MA induced Rac1 GTPase activity, while Rho GTPase inhibitor Y-27632 attenuated MA-induced endothelial permeability. We showed that benzene metabolites compromised the endothelial barrier by altering HSP- and GTPase-related signaling pathways.

## Introduction

Benzene is an airborne toxicant, abundant in both indoor and outdoor air. It is one of the top twenty chemicals emitted by industrial sources in the United States (1) and is ranked sixth on the Agency of Toxic Substances and Disease Registry’s substance priority list. Prominent sources of atmospheric benzene include petrochemicals, combustion of organic chemicals, tobacco products, and household items, including personal care products, cleaning products, paints, furniture wax, etc. (1). Levels of benzene in the ambient air <100 ppb; however, the atmospheric concentration of benzene can exceed 50 ppm near the emission source. The U.S. Occupational Safety and Health Administration has established an occupational benzene exposure limit of 1 ppm; however, occupational benzene exposure can exceed 100 ppm in developing countries (2).

Benzene is a well-known carcinogen and hematopoietic toxin (1), and emerging data suggest that benzene exposure is associated with insulin resistance (3), diabetes (4), and cardiopulmonary diseases (5–10). However, the mechanisms of benzene-induced cardiovascular disease remain unknown. We observed that benzene exposure affects endothelial cells and endothelial progenitor cells associated with endothelial repair (10). Our studies with benzene exposure in mice subjected to pressure overload-induced cardiac dysfunction (6) shows increased lymphocyte extravasation into cardiac tissues, indicating potential changes in endothelial permeability, lymphocyte trafficking, or both.

Alteration of microvascular endothelial permeability by inflammation or due to mechanical or chemical stress can induce myocardial interstitial edema, which plays an essential role in the pathology of heart failure (11, 12). Accumulation of interstitial fluid impaired myocyte contractile function, induced fibrosis, and compromised cardiac output (13, 14). Moreover, heart failure with preserved ejection fraction is increasing in prevalence in the aging and diseased human population, but the mechanisms responsible for disease manifestation are not well understood. Microvascular leakage and endothelial dysfunction contribute to myocardial fibrosis and diastolic heart failure (15). Therefore, changes in cardiac microvascular permeability can markedly affect heart function, especially under pathophysiological conditions.

Because most of the benzene toxicity is exerted by its active metabolites, this study examined the effect of benzene metabolite exposure on vascular permeability and delineated the underlying molecular mechanisms.

## MATERIALS AND METHODS

### Reagents

Benzene permeation tubes were obtained from Kin-Tek (La Marque, TX). Primers and probes for real-time PCR were purchased from Integrated DNA Technologies (Coralville, IA) and ThermoFisher Scientific (Waltham, MA). All other chemicals and enzymes were from Sigma Chemical Co. (St. Louis, MO) or Invitrogen (Carlsbad, CA). Benzene metabolite trans,trans-mucondialdehyde (MA) was prepared from muconic acid (Sigma-Aldrich, St. Louis, MO) as described previously (16). The structure and purity of MA were established by proton and carbon NMR and melting point.

### Animal housing and maintenance

C57BL6/J 6-12 weeks old male mice obtained from Jackson Laboratory (Bar Harbor, ME) were maintained on standard chow in a pathogen-free facility accredited by the Association for Assessment and Accreditation of Laboratory Animal Care. The University of Louisville Institutional Animal Care and Use Committee approved all procedures. We used male mice in our study because multiple studies have shown that the rate of benzene metabolism, and consequently benzene toxicity, is significantly higher in male mice than female mice (17–19).

### Cell culture

Mouse cardiac microvascular endothelial cells (CMVEC) and human aortic endothelial cells (HAEC) were purchased from CellBiologics, Chicago, IL. Cells were cultured in a complete endothelial growth medium without phenol red (CellBiologics) and used up to 8 passages for the experiments.

### Microparticle measurement

Adult male C57BL/J6 mice were administered MA (1 mg/kg body weight, dissolved in 5% DMSO in PBS) or the vehicle by retroorbital injection. The mice were euthanized 24 hours later (by pentobarbital, i.p.), and the peripheral blood was collected by heart puncture. Microparticles in the peripheral blood were measured as described before (20, 21). Briefly, the plasma was centrifuged for 2 min (11,000 ×g at 4 °C) to remove residual cells and debris, and the supernatant was aspirated and centrifuged for 45 min (17,000 ×g at 4 °C). The resulting microparticle pellet was resuspended in Annexin V Buffer pre-filtered through 0.22 μm syringe filter and incubated with the anti-mouse FcBlock (CD32/CD16) for 10 min. Microparticle (<1 μm in size and positive for Annexin V staining) subpopulations were identified based on expression of various surface markers (endothelial microparticles-<1 μm/AV+/Flk+, activated endothelial microparticles-<1 μm/AV+/CD62E+, endothelial progenitor cells microparticles-<1 μm/AV+/Flk+/Sca+. Identical samples with no antibodies were utilized as controls for the gating. Counting beads added to individual samples were used for data normalization.

### Endothelial permeability *in vivo*

Vascular permeability *in vivo* was performed as described (22). Briefly, adult male C57BL6/J mice were anesthetized with isoflurane, and histamine inhibitor pyrilamine maleate was intraperitoneally administered at a dose of 40 µg per 1 gram of body weight to suppress histamine release at the intradermal injection site. Next, the skin around the belly was shaved, and mice were injected intravenously with 100 µL of 1% Evans blue dye. Thirty minutes later, the mice were injected with MA or HQ (400 pmoles in 20 µL of 5% DMSO) per injection site. Mice treated with the vehicle served as controls. After 30 min, the skin around the injection sites was dissected and dried overnight at 55 °C. The skin weight was measured for subsequent normalization, and the dye was then eluted from the dissected samples with formamide at 55 °C. Following centrifugation at 10,000 g for 40 min, the optical density of the supernatant was measured at 620 nm.

### Measurements of transendothelial monolayer resistance

Transmonolayer electrical resistance of CMVEC grown on gold electrodes in NSP-96 CardioExcyte96 Sensor Plates (Nanion Technologies, Livingston, NJ) with standard 2 mm recording electrode was measured with the electrical cell impedance sensor technique using the CardioExcyte96 impedance sensing system (Nanion Technologies, Livingston, NJ). Endothelial cells were seeded at 2.5 × 10^4^ cells/well density and incubated for 48 hours at 37°C. Impedance was recorded after cell attachment and continued until cells became confluent and the increase in impedance plateaued. The impedance data were normalized to the initial resistance and plotted as normalized impedance.

### RNA Isolation and RNAseq analysis

HAEC were incubated with MA (10 µM in 0.1% DMSO) or vehicle (0.1% DMSO) for 2, 6, and 24 hours, and total RNA was isolated using an RNeasy Mini Kit (Qiagen). RNA quality was measured by Agilent 2100 bioanalyzer (Thermo Fisher Scientific, MA, USA), and samples with high RNA integrity were used for subsequent RNAseq analysis. RNA samples were processed by Novogene using mRNA sequencing services (Novogene, Beijing, China). The resultant raw reads of the FASTQ files were aligned to the human genome (hg38) using the HISAT2 R package (23). The mRNA differentially regulated genes (DEG) and pathway enrichment analysis were performed using DESeq2 (24) and ReactomePA enrichPathway (25) R packages.

### Endothelial permeability *in vitro*

Endothelial permeability assay was performed using cell culture inserts as described previously (26). The transwell PC membranes (1.0 μm pore, 24 well, membrane 33 mm^2^) were coated with rat collagen type I (CellBiologics) at a 7.5 µg/cm^2^ concentration. CMVEC were seeded onto membranes at a density of 1.0 × 10^5^ cells per membrane. Cells on transwell membranes were maintained at 37 °C in a CO_2_ incubator for 2-3 days to reach confluence. To assess cellular monolayer permeability, Alexa 488 cadaverine (MW 0.64 kDa), Texas Red Dextran (MW 70 kDa), and FITC Dextran (MW 500 kDa) from ThermoFisher were added to the luminal chamber at a concentration of 1 µM in 200 µL of growth media without FBS and phenol red. The fluorescence intensity in the lower chamber was measured using a Synergy H1 fluorescence microplate reader (Biotek, USA) at 1, 2, 3, 4, 5, 6, 7, and 8 hours after adding fluorescent dextrans. After fluorescence measurement, the media was returned to the lower chamber. MA was added to the media at the final concentration of 10 µM to determine its effect on endothelial permeability. The effect of HSP and GTPase inhibitors was analyzed by adding allosteric HSP70 inhibitor JG98 at a final concentration of 2 µM 60 min before, HSP90 inhibitor 17-AAG at a final concentration of 2 µM 2 hours before, HSP70 inducer TRC051384 at final concentration 10 µM 4 hours before, selective ROCK1 (p160ROCK) inhibitor Y-27632 at final concentration 10 µM, 30 min before exposure to MA. The concentration of fluorescent dextranes in the lower chamber was calculated using a calibration curve with known dextran concentrations.

### Measuring Rho1 GTPase activity

The Rac1 GTPase activities were measured by G-LISA Rac1 activation assay kit (Cytoskeleton Inc., Denver, CO) in cell lysates prepared from CMVEC exposed to 10 µM MA for 6 hours. Briefly, CMVEC were lysed at the indicated time points using the provided cell lysis buffer and protease inhibitor cocktail. Protein concentration in lysates was quantified using the precision red protein assay. The exact amount of protein lysate was loaded into each well, and the signal was read by measuring absorbance at 490 nm using a microplate reader.

### Statistical Analyses

Data are presented as means ± SEM. The statistical significance of differences was determined by t-test. A two-way analysis of variance (ANOVA) with Bonferroni *post hoc* test was used for time-dependent dextran diffusion data analysis. One one-way ANOVA with Tukey *post hoc* test was used to compare differences between multiple treatment groups. A p-value of <0.05 indicated statistically significant differences. All analyses were performed using Excel and GraphPad Prism software (GraphPad Software, San Diego, CA).

## RESULTS

### Effect of benzene metabolites on endothelial cell permeability *in vitro*

Most of the toxicity of benzene is exerted by its active metabolites. Therefore, we examined the effect of benzene metabolites – hydroquinone, catechol, and t,t-muconaldehyde (MA) on the impedance of cardiac microvascular endothelial cells (CMVEC). As shown in **Fig. 1A and Fig. 1B**, MA decreased impedance in a time- and dose-dependent manner, whereas hydroquinone (10 µM) and catechol (10 µM) did not affect CMVEC impedance. Furthermore, MA increased endothelial permeability to 70 kDa dextran (**Fig. 1C**) and 500 kDa (**Fig. 1D**) but did not affect the permeation of low molecular mass molecule spermidine (**Fig. 1E**).

**Figure 1.**
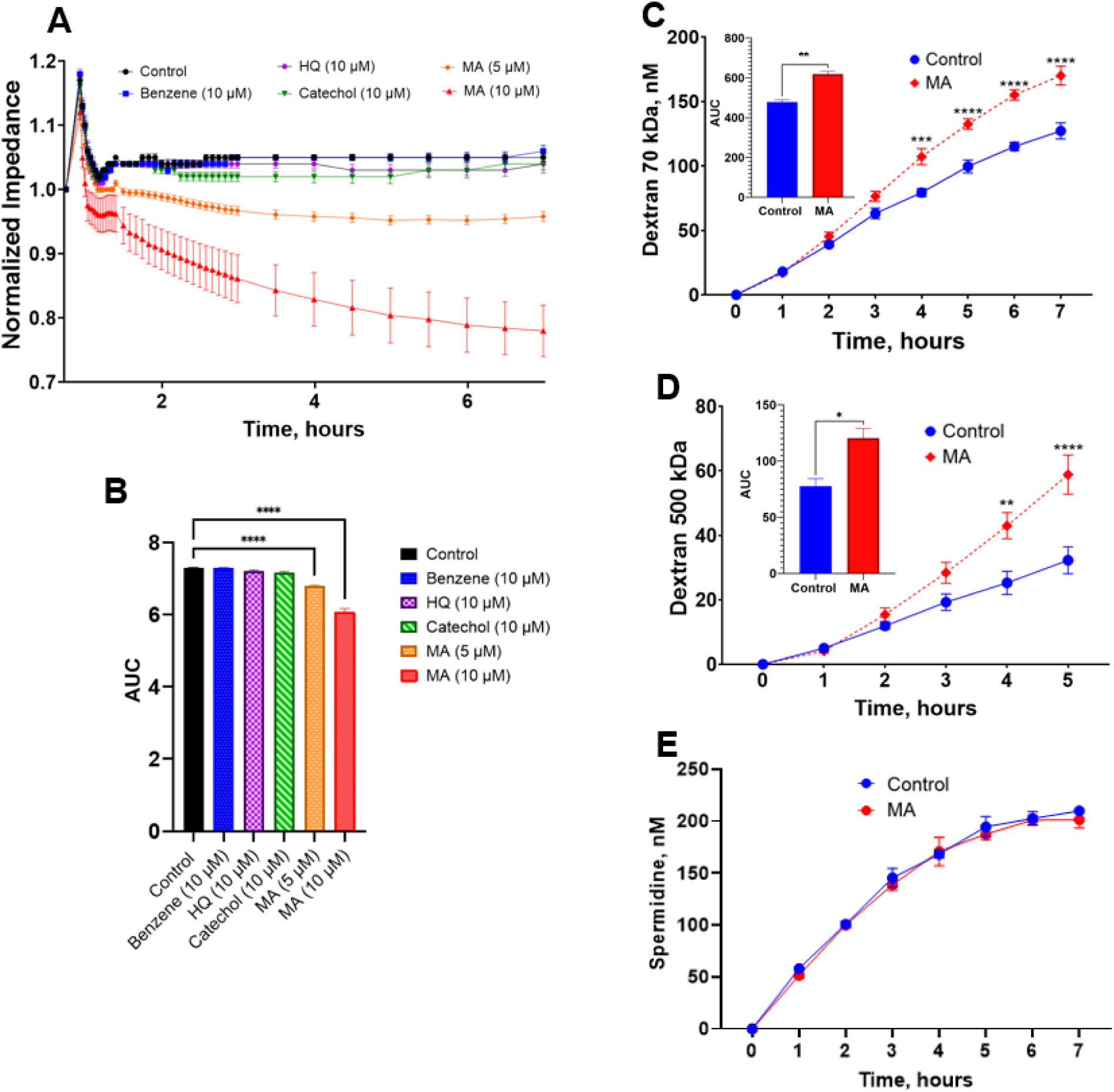
Benzene metabolites increase vascular permeability in vitro. **A,** The normalized impedance of mouse cardiac microvascular endothelial cells plated on microelectrodes and exposed to benzene and its metabolites. Electrical resistance across the endothelial monolayer was measured using the electrical cell-substrate impedance sensing system for 7 hours as described in Materials and Methods. Data were normalized to impedances before the addition of benzene metabolites. The absolute baseline impedance values of respective endothelial cell layers were around 250 Ohm before metabolites addition. **B**, Area under the curve (AUC) for normalized impedance. Data are mean ± SE. ****P<0.0001, One-way ANOVA with Bonferroni post-test. **C-E,** The permeability of CMVEC monolayer to fluorescent dextrans with molecular weight 70 kDa (**C**), 500 kDa (**D**), and 0.1 kDa spermidine (**E**) in response to MA treatment. Data presented as mean ± SE. **P<0.01, ***P<0.001, ****P<0.0001, Two-way ANOVA with Tukey post test. The insets show the AUC. Data presented as mean ± SEM *P<0.01 versus controls following a two-tailed t-test (unpaired).

### Muconaldehyde induces endothelial permeability in mice

To examine the effect of MA on vascular permeability *in vivo*, adult male C57BL/6J mice were injected with 400 pmoles MA intradermally, and Evans Blue dye leakage was measured near the injection site. As shown in **Fig. 2A**, MA increased the vascular leakage by 54% (p<0.0001). Complimentary experiments with mice exposed to MA resulted in increased formation of endothelial microparticles and activated endothelial microparticles in the blood (**Fig. 2B**). These data suggest that MA compromises endothelial integrity and renders vasculature permeable to the plasma macromolecules.

**Figure 2.**
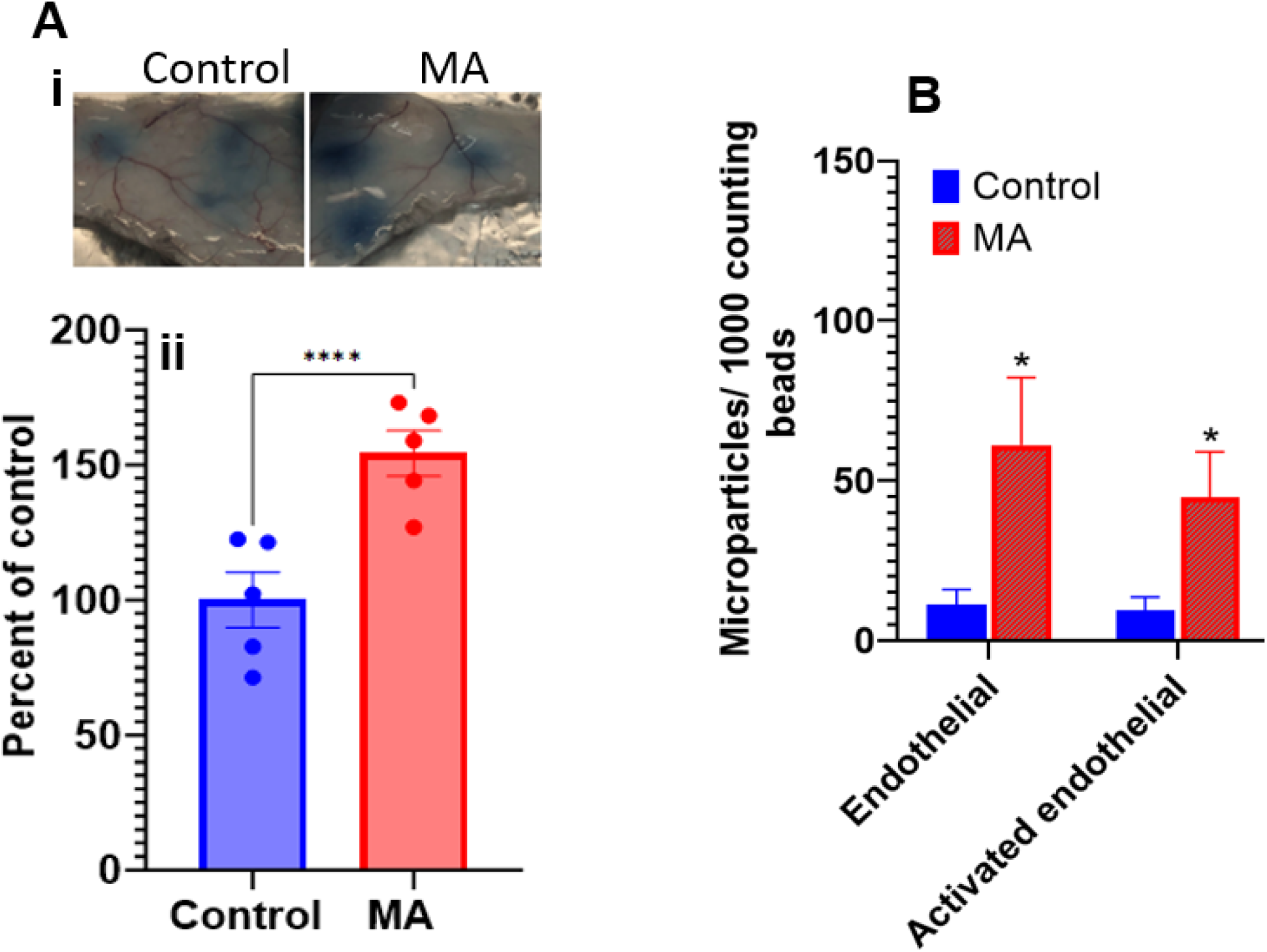
Muconaldehyde-induced endothelial permeability in mice.. **A.** Eight-week-old male C57BL/6 mice were injected (i.v.) with Evans blue and then treated with intradermal injections of MA (400 pmoles per injection) or the vehicle. (i) A representative photomicrograph of Evans blue leakage was taken at the site of subcutaneous injections 30 minutes after MA treatment. (ii) Group data of vascular leakage in the skin extracts 30 min after MA treatment. Data are mean ± SEM. ****P<0.0001 versus vehicle-treated control mice as per two-tailed t-test (unpaired). **B**. Endothelial and activated endothelial microparticles in the peripheral blood of MA-treated mice. Mice were injected (i.v.) with MA (1 mg/kg body weight) or vehicle, and endothelial microparticles were measured in the plasma using flow cytometry 24h after injection. Values are mean ± SEM. *P<0.05 versus vehicle-treated control mice as per two-tailed t-test (unpaired).

### Molecular mechanisms of muconaldehyde-induced endothelial permeability

To delineate the molecular mechanism by which MA worsens the endothelial cell integrity, we performed deep RNA sequencing. For these experiments, we used HAEC because increased endothelial permeability would promote leukocyte recruitment, vascular inflammation, and atherogenesis. The time course experiments revealed that MA differentially regulated the transcription of 224 unique genes at 2 hours, 965 genes at 6 hours, and 172 genes at 24 hours (padj < 0.05 and 2 < Log2(Fold Change) < -2; **Fig. 3A and Supplemental Table 1**). Molecular function enrichment analyses of differentially expressed genes as depicted by Reactome analysis showed significant enrichment of the genes associated with the regulation of *heat shock protein (HSP) binding, misfolded protein binding,* and *“DNA-binding transcription activator activity* after 2- and 6 hours of MA exposure (**Fig. 3Bi**). Specifically, MA robustly induced members of *Hsp70* (*Hspa6*, *Hspa7*, *Hspa1a*, *Hspa1b*) and to a lesser extent *Hsp90* (*Hsp90aa1*, *Hsp90b1*, *Hsp90ab1*, *Hsp90b2p*) family at 2- and 6 hours (**Fig. 3Bii**). Network analysis revealed that induction of HSP70 genes is associated with the activation of *cellular response to stress*, *HSF-1-dependent transactivation*, and *attenuation phase* pathways (**Fig. 3Biii**). MA exposure also down-regulated the genes associated with CXCR chemokine receptor binding (2h), GTPase regulator activity and nucleotide-triphosphate regulator activity (6h), and microtubule binding and tubulin binding (24 h). The induction of HSPA1B by MA was confirmed at mRNA and protein levels (**Fig. 3C**). Similar to HAEC, MA also increased levels of the HSPA1B protein in CMVEC (**Fig. 3D**). The newly synthesized HSP70 colocalizes with F-actin foci, suggesting an interaction of HSP with the remodeling cytoskeleton (**Fig. 3E**) in MA-treated murine heart microvascular endothelial cells.

**Figure 3.**
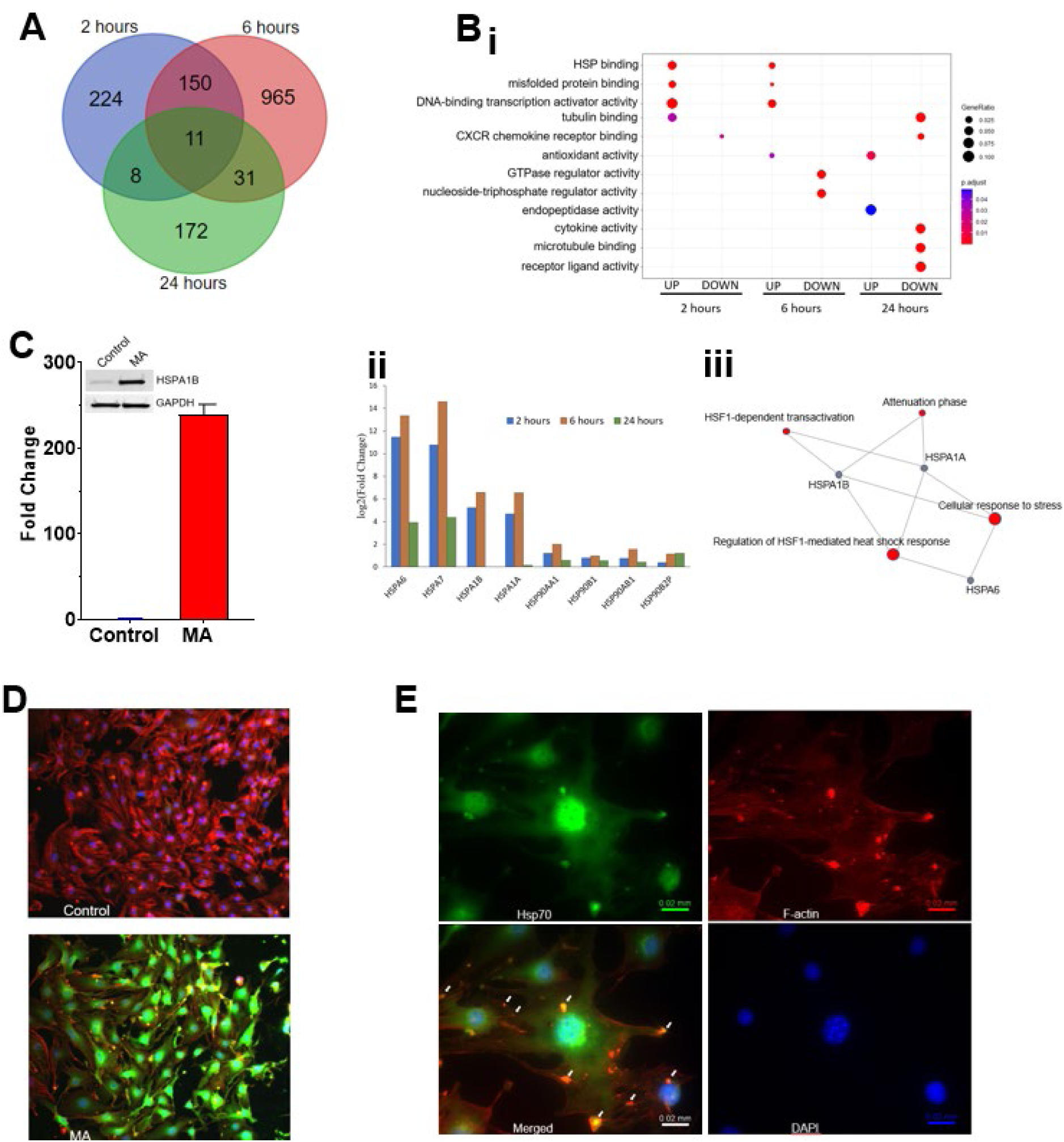
Muconaldehyde-induced stimulation of heat shock proteins in endothelial cells. Human aortic endothelial cells (HAEC) were incubated with MA (10 µM) for 2, 6, or 24h, and gene transcription was measured by RNA sequencing. **A,** Venn diagram of differentially expressed genes at indicated time points (padj < 0.05; log2FoldChange ≥ |2|). **B. (i)** Pathway enrichment analysis of MA-treated HAEC. (**ii**) HSP70 and HSP90 induction in MA-treated HAEC. (**iii)** Gene network of HSP-related pathways activated after 6h exposure to MA. **C,** Induction of HSPA1B in HAEC following exposure to 10 μM MA. **D,** Hspa1b induction in MA (10 µM) -treated CMVEC. **E,** Immunofluorescent staining of Hsp70 and F-actin in CMVEC treated with MA (10 µM) for 24h.

### Heat shock protein regulated muconaldehyde-induced endothelial permeability

To examine the contribution of HSPs in decreasing endothelial integrity, we examined the pharmacological inhibitor of HSP70s (17AAG) and HSP90s (JG98) on endothelial permeability of murine heart microvascular endothelial cells. As shown in **Fig. 4A-D**, both 17AAG and JG98 prevented MA-induced endothelial permeability (**Fig. 4A-D**). Conversely, HSP70 activator TRC051384 (TRC) increased the transmigration of 70 kDa dextran through endothelial cell monolayer (**Fig. 4E-F**). These data suggest that MA-induced HSPs compromise endothelial monolayer integrity and increase its permeability for macromolecules.

**Figure 4.**
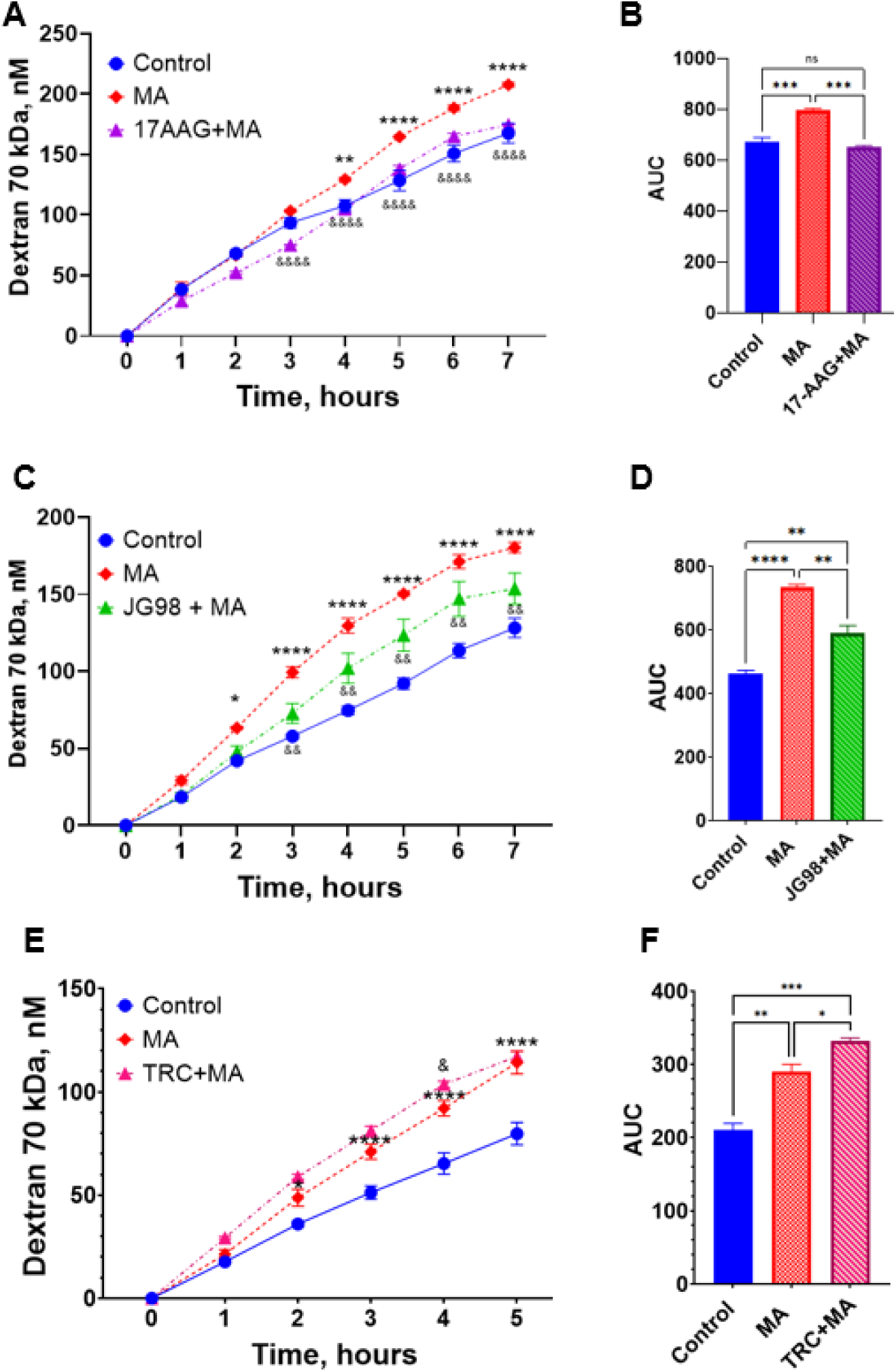
Permeability of microvascular endothelial cells in response to HSP inhibitors and activators. The endothelial monolayer was exposed to the medium with 0.1% DMSO in PBS or 10 µM MA for indicated times. Fluorescent dextran concentrations were measured in lower chambers as described under *Material and Methods*. Cells were co-exposed to 2 μM the HSP90 inhibitor 17-AAG (**A**), 2 μM the HSP70 inhibitor JG98 (**C**) or 10 μM the HSP70 activator TRC (**E**). Data are shown as mean ± SE. *P<0.05, **P<0.01, ****P<0.0001 for MA vs. Control; ^&^P<0.05, ^&&^P<0.01, ^&&&&^P<0.0001 for inhibitor/activator/MA vs. MA, Two-way ANOVA with Tukey post-test. The area under the curve (AUC) was calculated for the effects of 17-AAG (**B**), JG98 (**D**), and TRC (**F**) on MA-induced permeability. Data are shown as mean ± SE. *P<0.05, **P<0.01, ***P<0.001, ****P<0.0001, One-way ANOVA with Bonferroni post-test.

### Contribution of Rho GTPase on MA-induced endothelial permeability

Because 6 hours of MA exposure affected the GTPase regulatory network and GTPases regulate cell cytoskeleton and tight junctions, we examined these pathways’ molecular regulation. Pathway enrichment analyses suggested that MA primarily affected the genes associated with RHO GTPase cycle, RHOA GTPase cycle, and CD42 GTPase cycle (**Fig. 5A**). Gene network associated with GTPase pathways indicated altered expression of multiple guanine nucleotide exchange factors (GEFs) and GTPase-activating proteins GAPs) (**Fig. 5B, C**). To examine the role of Rho kinase in affecting MA-induced endothelial cell permeability, we probed how inhibition of this pathway affects MA-induced macromolecule permeability. As shown in **Fig. 5D-E**, Rho-associated coiled-coil-containing protein kinase (ROCK) inhibitor, Y-27632 attenuated MA-induced permeability to 70 kDa dextran in murine heart microvascular endothelial cells, and this was accompanied by a dose-dependent MA-induced activation of Rac1 GTPase (**Fig. 5F**). Collectively, these data indicated that MA-induced dysregulation of Rho GTPase activity affects endothelial permeability.

**Figure 5.**
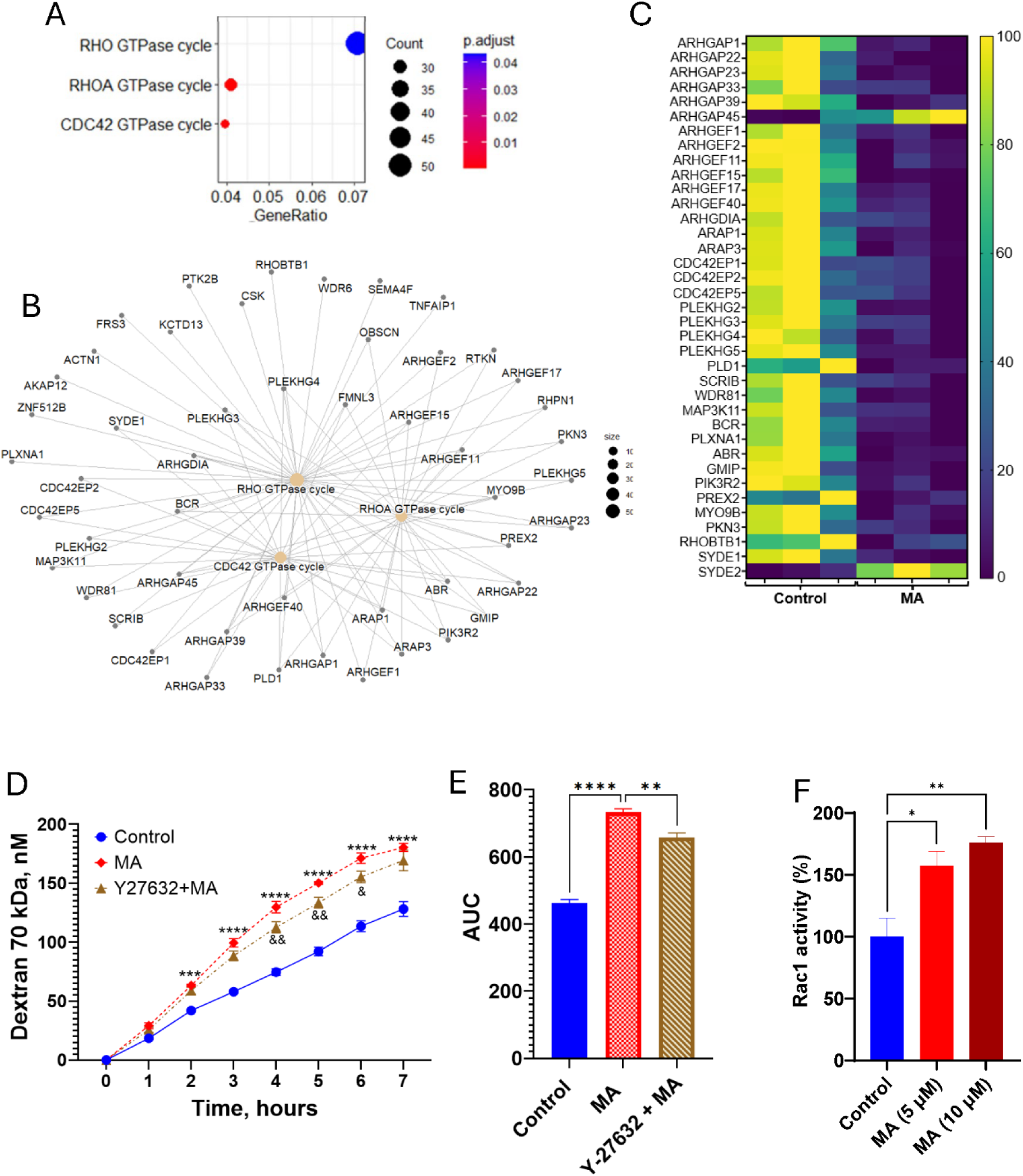
Effect of muconaldehyde on endothelial Rho GTPases. **A.** Dysregulated GTPase signaling pathways in HAEC incubated with MA(10 µM) for 6h. **B.** Gene-network for affected signaling pathways in MA-treated HAEC (6h). **C.** Heatmap of RhoA GTPase cofactors in MA-treated HAEC (6h). **D.** Effect of Rho-associated, coiled-coil containing protein kinase (ROCK) inhibitor (10 µM) on MA-induced permeability of MCMVEC. Data are mean ± SEM. * P<0.05 versus controls following two-way ANOVA with Tukey post test. **E.** Area under the curve (AUC) for 70 kDa dextran kinetic in MA-treated MCMVEC following one-way ANOVA with Tukey post-test. **F.** Effect of MA (10 µM, 6h) on Rac1 activity in MCMVEC. Data are mean ± SEM. *P<0.05 following one-way ANOVA with Tukey post-test.

## DISCUSSION

Our studies reveal that benzene metabolites compromise the microvascular endothelial barrier and cause vascular leakage and endothelial damage both *in vivo* and *in vitro*. These findings corroborate arsenic (27), trichloroethylene (28), formaldehyde (29), organochlorine pesticide endosulfan (30) and cigarette smoke (31)-induced increase in endothelial permeability. Our recent studies have shown that benzene exposure enhances the accumulation of leukocytes in the myocardium and compromises cardiac function in mice subjected to pressure overload (6). Epidemiological studies suggested that benzene exposure is associated with an increased risk of heart failure (9), ischemic stroke(32), type 2 diabetes (33), and dose-dependent increase in mortality risk from all-cause cardiovascular disease (34). Because compromised vascular permeability and endothelial integrity promote cardiovascular disease, it is plausible that a benzene-induced increase in the leukocyte trafficking in the failing heart is facilitated by endothelial toxicity and vascular leakage by its active metabolites.

In mammals, benzene is readily metabolized to phenol, catechol, hydroquinone, and MA (35, 36). Being an unsaturated aldehyde, MA is highly reactive and can induce hematopoietic toxicity (36, 37), promote endothelial cell apoptosis (8), and inhibit gap junction intercellular communication (38, 39). Gap junctional intercellular communication between adjacent cells typically facilitates the exchange of low molecular weight molecules (up to 1.2 kDa) (40) and enables cell-to-cell communication. Our data demonstrate increased endothelial permeability in response to MA exposure, suggesting disruption of tight and adherent junctions organization and integrity. These observations indicate that MA exerts a different mode of action on tight and adherent junctions than on gap junctions.

HSPs are an evolutionarily conserved group of proteins (classified by their molecular weights) that serve as molecular chaperones and help to keep the proteins in their native structures. Increased expression of HSP by MA corroborates short-term ischemia-induced upregulation of HSP70 in the rabbit heart (41). A recent study has also reported the expression of HSP70 members - *Hspa1a* and *Hspa1b* genes in murine atherosclerotic plaque macrophages (42). *In vitro*, HSP expression also increases in response to atherogenic stimuli, including oxidized LDL, oxidative stress, mechanical stress, inflammation, infection, and increased temperature (43–45). Induction of HSPs enhances protein folding, inhibits inflammation and apoptosis, provides cytoskeletal protection, and increases nitric oxide synthesis (46). HSP90 plays a critical role in multiple pathways linked to cardiac dysfunction, including ventricular hypertrophy and associated fibrosis (47, 48). Moreover, the pharmacological inhibition of HSP90 prevented myocardial ischemia and pressure overload-induced cardiac dysfunction (47, 49). Inhibition of HSP70 attenuated angiogenesis in a phosphatidylinositol 3-kinase/Akt pathway-dependent manner (50). Our observation that HSP70 and HSP90 inhibitors attenuated MA-induced endothelial permeability while HSP70 activator TRC augmented MA-induced endothelial permeability suggests that HSPs facilitate benzene-induced vascular leakage and compromised endothelial integrity. Our studies are also consistent with the observations that HSP90 inhibition prevents TGF-β1-induced pulmonary endothelial cell permeability *in vitro* (52) and endotoxin-induced capillary leak and acute lung injury in mice (53) and protects blood-brain barrier integrity in cerebral ischemic stroke (54). Conversely, a recent study by *Yuan et al.* indicated that Hsp70 chaperones can act as a positive regulator of endothelial barrier function in LPS-induced acute lung injury (51); however, in this model, the administration of LPS decreased levels of HSP70 expression while chemical inducer of HSP70 attenuated vascular leakage. Therefore, it is likely that HSP70 might have a protective or disruptive effect on endothelial monolayer permeability depending on the location, type of vessel, and nature of the stressor.

The molecular mechanisms responsible for the protective effect of HSP90 inhibitors observed in our study are unclear. However, it might be attributed to the attenuation of RhoA function, which prevents the cytoskeleton’s remodeling (55, 56). Recent studies indicated that HSP90 can participate in the organization and regulation of cytoskeleton in endothelial and epithelial cells. HSP90 binds F-actin (70, 71) and Wiskott-Aldrich syndrome protein (N-WASP) (72) and is directly involved in actin filament assembly. The destabilization of actin microfilament assembly increases endothelial monolayer permeability in different vascular beds (57, 58). HSP90 also protects LIM kinase1 degradation and enables actin polymerization through modulation of p-cofilin levels (59). Alternatively, GTPase activity could be regulated by indirect modification of upstream guanine nucleotide exchange factors that regulate the activity of Rho and, subsequently, the function of Rho-associated kinases ROCK1 and ROCK2 (60). Rac1, RhoA, and Cdc42 represent the Rho subfamily of GTPases and regulate a wide range of cellular processes, including cell-cell interaction, cell adhesion, and cytoskeleton reorganization (61). The activity of Rho GTPases is regulated by a complex network of guanine nucleotide exchange factors (GEFs), GTPase-activating proteins (GAPs), and guanine nucleotide dissociation inhibitors (GDIs), which determine the activation or inhibition of specific GTPases, and hence facilitate a particular cellular outcome (62). GTPases play a critical role in the maintenance of endothelial barriers and in controlling extravasation (63). Rac1 plays a role in maintaining endothelial barrier integrity through actin cytoskeleton strengthening and an increase in the stiffness of the cell peripheral membrane under inflammatory conditions (64). Our results showing the attenuation of MA-induced endothelial permeability by Y-27632 inhibitor of ROCK1 and ROCK2 support the importance of Rho GTPase in this process and are in agreement with previously published protective effects of Y-27632 against thrombin-induced endothelial barrier failure (65–67).

Collectively, our data suggest that reactive benzene metabolites can damage the endothelium and disrupt the homeostasis of the microvascular endothelial monolayer, making it permeable to solutes and macromolecules. These processes could be regulated by HSPs and Rho GTPase.

## Acknowledgement

**This study was partially supported** by NIH grants R21 ES033323, P42 ES023716, R01 HL149351, R01 HL137229, R01 HL146134, R01 HL156362, R01 HL138992, R01 HL122676, U54 HL120163, and the Jewish Heritage Foundation grant OGMN190574L.

